# Nextstrain: real-time tracking of pathogen evolution

**DOI:** 10.1101/224048

**Authors:** James Hadfield, Colin Megill, Sidney M. Bell, John Huddleston, Barney Potter, Charlton Callender, Pavel Sagulenko, Trevor Bedford, Richard A. Neher

## Abstract

**Summary:** Understanding the spread and evolution of pathogens is important for effective public health measures and surveillance. Nextstrain consists of a database of viral genomes, a bioinformatics pipeline for phylodynamics analysis, and an interactive visualisation platform. Together these present a real-time view into the evolution and spread of a range of viral pathogens of high public health importance. The visualization integrates sequence data with other data types such as geographic information, serology, or host species. Nextstrain compiles our current understanding into a single accessible location, publicly available for use by health professionals, epidemiologists, virologists and the public alike.

**Availability and implementation:** All code (predominantly JavaScript and Python) is freely available from github.com/nextstrain and the web-application is available at nextstrain.org.

---

Viral pathogens pose an ongoing and ever-present danger to global human health, with recent epidemics such as the West-African Ebola epidemic and the ongoing Zika epidemic in the Americas highlighting the risk. The rapid evolution of these viruses allows inference of epidemic history from genomic data. Such analyses are often done in isolation, and may lack the spatial or temporal context in which to best interpret the results, which impedes understanding [1]. Furthermore, the results of analyses are often not made available to the public or health bodies until after publication, which may be too late to effect change in policy or understanding. We have developed Nextstrain to visualise outbreaks in as close to real time as possible. Whilst currently encompassing a selection of viruses, extension to non-viral pathogens is forthcoming.

The regularly updated nature and rapidity of these analyses is crucial to the monitoring and understanding of pathogen epidemiology and evolution. Sequencing times and costs are continually dropping, with on-the-ground sequencing being deployed during the recent West African Ebola and Zika epidemics [2, 3]. However, this speed of sequencing must be complemented with rapid methods by which to analyse, interpret, and disseminate results.

Nextstrain consists of three components: *Fauna* is a collection of Python scripts to maintain a database representative of all available sequences and related metadata, sourced from public repositories such as NCBI, GISAID and ViPR, as well as GitHub repositories and other sources of genomic data. *Augur* is the bioinformatics pipeline, performing subsampling, alignment, phylogenetic inference, temporal dating of ancestral nodes and discrete trait geographic reconstruction, including inference of the most likely transmission events. This leverages the maximum likelihood phylodynamic analyses implemented in TreeTime [4], allowing a full analysis of the entire Ebola epidemic (n=1581 genomes) in under 2 hours on a modern laptop computer. These scripts separate generic core functionality from a light pathogen-specific layer such that they are easily co-opted to different pathogens. Outputs from the pipeline are uploaded online and immediately available through the visualisation software *Auspice*, located at nextstrain.org. This approach is similar in concept to Nextflu [5] however extended and generalised to different viral pathogens. There is a growing need for surveillance of non-influenza viruses [6], and Nextstrain is able to be extended to nearly all outbreaks with readily accessible genomic data.

## Joint temporal and spatial visualisation

Conveying understanding of pathogen evolution through space and time involves filtering large amounts of data into forms that can be easily reasoned with. Auspice employs linked views into different facets of the data, whereby changes in one view are reflected in all others. These views allow simultaneous interrogation of phylogenetic and spatial relationships, with additional data such as genotype or serotype expressed through colourings (Figure 1). This is coupled with an interactive time slider to see how the pathogen has evolved and spread over the course of the epidemic. By animating the temporal dimension, a high level overview of how the entire outbreak unfolded is quickly gained. This approach both communicates the geographical spread of the epidemic alongside the underlying genomic data that supports this geographic reconstruction.

**Figure 1.**
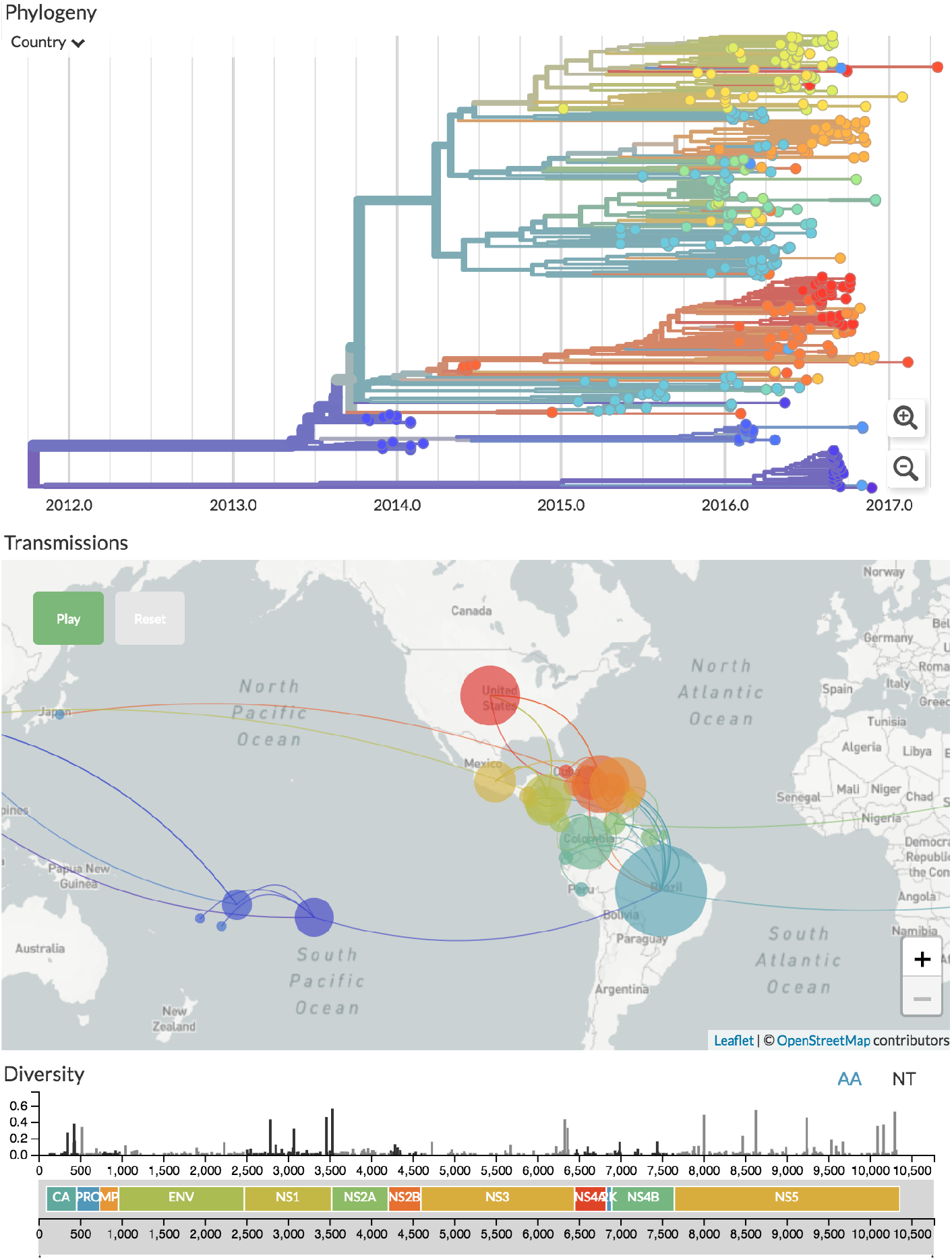
Genomic epidemiology of Zika virus as of Oct 2017 (live display at nextstrain.org/zika). The main interface consists of three linked panels — a phylogenetic tree, geographic transmissions, and the genetic diversity across the genome.

Maximum likelihood ancestral state reconstruction of discrete traits such as country or region of isolation allows identification of probable transmission events given the sampled data, together with a marginal likelihood for each event. Inferred transmissions are displayed as colours on the tree and each transmission is also drawn on the map. It is important to make known the confidence of any such reconstruction, and we do this in two ways: by matching colour saturation to the confidence of that trait, and by displaying all relevant information when one hovers over the corresponding branch or isolate on the tree.

## Monitoring of evolution and adaptation

Nextstrain tracks and reconstructs mutations across the tree and displays this information as a bar-chart of entropy at each position in the genome, as well as showing the mutations inferred to occur on each branch by hovering over the tree. Selecting a position in the genome with non-zero entropy reveals the distribution of the segregating variant in the phylogeny and on the map. This allows interrogation of genetic change which may be adaptive or underlying a change in disease dynamics.

For many pathogens, the emergence and spread of gain-of-function variants is a grave concern. For instance, China has experienced seasonal epidemics of influenza A/H7N9 over the past five years, however human to human transmission is thought to be rare, with the majority of human cases being spillover from poultry. The threat of mutations which facilitate human to human transmission is of extreme concern, as H7N9 has a mortality rate of around 30% [7]. Nextstrain allows monitoring of experimentally determined mutations, such that interventions and control measures may be implemented where appropriate.

## A model for public sharing of data

Nextstrain currently presents a single, continuously updated overview of both endemic viral disease (seasonal influenza, dengue) as well as emergent viral outbreaks (avian influenza, Zika, Ebola), all based upon the same underlying bioinformatics architecture. This architecture is well positioned to respond to future outbreaks, be they viral or bacterial.

Analysis of such outbreaks relies on public sharing of data, and Nextstrain has the ability to automatically update as new sequences from a range of public databases and repositories appear. Scientists are justifiably hesitant to cede control of their data, and we try to address these concerns by preventing access to the raw genome sequences, and by clearly indicating the source of each sequence. Derived data, such as phylogenetic trees, metadata and screenshots may be downloded, and private metadata may be added by users by simply dragging a CSV file onto the browser. We believe this strikes a compromise between keeping certain data private and allowing the dissemination of results important to the wider scientific community, thereby encouraging collaboration between scientists. Genomic epidemiology has the potential to inform the public, health organisations and scientists alike, a potential realised by sharing of data in real-time rather than retrospectively [8].

